# Unpredictability of the fitness effects of antimicrobial resistance mutations across environments in *Escherichia coli*

**DOI:** 10.1101/2023.10.25.563906

**Authors:** Aaron Hinz, André Amado, Rees Kassen, Claudia Bank, Alex Wong

**Affiliations:** Department of Biology, Carleton University, Ottawa, ON K1S 5B6, Canada; Department of Biology, University of Ottawa, Ottawa, ON K1N 6N5, Canada; Department of Biology, McGill University, Montreal, QC H3A 1B1, Canada; Institute of Ecology and Evolution, University of Bern, Switzerland; Swiss Institute of Bioinformatics, Lausanne, Switzerland; Gulbenkian Science Institute, Oeiras, Portugal; Institute for Advancing Health Through Agriculture, Texas A&M System, Fort Worth, Texas, USA

**Keywords:** Antimicrobial Resistance Mutations, Costs of Resistance, Genotype-by-Environment Interactions, Epistasis, Fitness Landscapes, Rough Mount Fuji Model

## Abstract

The evolution of antimicrobial resistance (AMR) in bacteria is a major public health concern. When resistant bacteria are highly prevalent in microbial populations, antibiotic restriction protocols are often implemented to reduce their spread. These measures rely on the existence of deleterious fitness effects (i.e., costs) imposed by AMR mutations during growth in the absence of antibiotics. According to this assumption, resistant strains will be outcompeted by susceptible strains that do not pay the cost during the period of restriction. Hence, the success of a given intervention depends on the magnitude and direction of fitness effects of mutations, which can vary depending on the genetic and environmental context. However, the fitness effects of AMR mutations are generally studied in laboratory reference strains and estimated in a limited number of environments, usually a standard laboratory growth medium. In this study, we systematically measure how three sources of variation impact the fitness effects of AMR mutations: the type of resistance mutation, the genetic background of the host, and the growth environment. We demonstrate that while AMR mutations are generally costly in antibiotic-free environments, their fitness effects vary widely and depend on complex interactions between the AMR mutation, genetic background, and environment. We test the ability of the Rough Mount Fuji genotype-fitness model to reproduce the empirical data in simulation. We identify model parameters that reasonably capture the variation in fitness effects due to genetic variation. However, the model fails to accommodate variation when considering multiple growth environments. Overall, this study reveals a wealth of variation in the fitness effects of resistance mutations owing to genetic background and environmental conditions, that will ultimately impact their persistence in natural populations.

**Author’s Abstract:** The emergence and spread of antimicrobial resistance in bacterial populations poses a continuing threat to our ability to successfully treat bacterial infections. During exposure to antibiotics, resistant microbes outcompete susceptible ones, leading to increases in prevalence. This competitive advantage, however, can be reversed in antibiotic-free environments, due to deleterious fitness effects imposed by resistance determinants, a concept referred to as the ‘cost of resistance’. The extent of these fitness effects is an important factor governing the prevalence of resistance in natural populations. However, predicting the fitness effects of resistance mutations is challenging, since their magnitude can change depending on the genetic background in which the mutation arose and the environmental context. Comprehensive data on these sources of variation is lacking, and we address this gap by determining the fitness effects of resistance mutations introduced in a range of *Escherichia coli* clinical isolates, measured in different antibiotic-free environments. Our results reveal wide variation in the fitness effects, driven by irreducible interactions between resistance mutations, genetic backgrounds, and growth environments. We evaluate the performance of a fitness landscape model to reproduce the data in simulation, highlight its strengths and weaknesses, and call for improvements to accommodate these important sources of variation.

## Introduction

Pervasive antibiotic use selects for bacteria with antimicrobial resistance (AMR) and has led to rising prevalence of multidrug-resistant pathogens [1–4]. Considering the slow pace of antibiotic discovery [5], effective stewardship of existing antibiotics is essential. Antibiotic restriction is a widely used approach that aims to reverse the spread of resistance by reducing the selective pressure that maintains AMR in bacterial populations. Antibiotic restriction in medical, agricultural, and veterinary settings has correlated with decreased resistance, although responses vary widely, with resistance rarely eliminated, and some efforts failing entirely [6–10]. The ability to predict success or failure of antibiotic restriction will be important for implementing more rational interventions.

The premise behind antibiotic restriction is that resistant microbes are outcompeted by susceptible microbes in antibiotic-free environments due to fitness costs imposed by resistance determinants. These costs can derive from functional tradeoffs of altering antibiotic targets, unregulated expression of drug efflux pumps, or burdens of maintaining replicating resistance plasmids [11]. However, while evidence indicates that AMR mutations are generally costly [12], several mechanisms can promote AMR persistence even in the absence of direct antibiotic selection. First, resistance to a restricted antibiotic can be indirectly selected by the presence of non-restricted antibiotics via mechanisms of cross-resistance [13–15] or co-selection of genetically linked resistance determinants [16–18]. Second, some AMR mutations may incur little or no cost to the microbe, allowing resistance to be maintained in antibiotic-free environments [19]. Third, costs might be heterogeneous across different environments, allowing for resistance to be maintained in cost-free environmental refuges [20]. Finally, second-site compensatory mutations, either segregating in the population or arising after resistance evolution, can reduce fitness costs without loss of resistance [11].

Knowledge of the range of fitness effects caused by AMR mutations is crucial to guide decision-making but comprehensive data are lacking. Standard practice is to obtain experimental measures of the fitness effects of AMR mutations from well-characterized laboratory strains grown in standard growth media [12,21]; however, it has become increasingly evident that these fitness effects can be modulated by both genetic and environmental variation [22–25]. For example, the magnitude or the direction (costly vs. beneficial) of a mutation’s fitness effect may change depending on the genetic background in which the mutation evolved, a form of genotype by genotype (G x G) interaction known as epistasis [26,23,24]. The growth environment can also impact fitness effects in ways that are hard to anticipate, a form of genotype by environment (G x E) interaction [22,27]. Furthermore, higher order interactions between the nature of the AMR mutation itself (modification of a target site versus deregulation of an efflux pump, for example), the genetic background on which the mutation occurs, and the growth environment (G x G x E interactions) can further complicate matters, potentially undermining predictions based on data from single genotypes and environments [28,22,29]. Currently we know very little about the extent to which the fitness effects of AMR mutations are consistent or variable across genotypes and environments. Obtaining such data is an important step towards predicting the success of antibiotic restriction strategies.

Ultimately, it will never be possible to empirically measure fitness for every mutation-genotype combination in all environments a microbial strain could encounter. Theoretical modeling could offer a complementary approach to predict the fitness of microorganisms across the various environments they populate. Two different types of models provide a rough prediction of antibiotic resistance fitness landscapes. Hill curve models predict fitness across antibiotic gradients but disregard other sources of environmental variation (see [30]). Alternatively, approaches based on Fisher’s Geometric Model tend to have more general applicability, but have many parameters, require extensive datasets, and are in practice cumbersome to fit [31–33]. Probabilistic fitness landscape models, such as the Rough Mount Fuji (RMF) model [34], are a third type of model that capture the relationship between genotype and fitness. These models are appealing because they feature tunable epistasis and are determined by few parameters [35]. In addition, they reasonably approximate some experimental fitness landscapes (e.g., [36]).

In this study, we present an empirical analysis of genetic and environmental factors that contribute to the variation in fitness effects among resistance mutations in *Escherichia coli* and evaluate the performance of an RMF-based genotype-fitness model to reproduce the empirical results. We introduced 7 resistance mutations individually into each of 12, primarily clinical, strains and quantified each resistance mutation’s fitness effect in four distinct growth environments. Overall, we show that fitness effects are extensively modulated by all three sources of variation: the type of AMR mutation, the genetic background, and the growth environment. The RMF model, despite having few parameters, reproduced single environment empirical results well but was unable to recover their full complexity when all environments were considered together. Our study highlights the challenges of predicting fitness effects from empirical data obtained from limited genetic backgrounds and environments while at the same time calling for improved fitness landscape models that account for these important sources of variation.

## Results

### Library of *E. coli* clinical isolates with introduced resistance mutations

We constructed a factorial mutant library by introducing each of 7 antimicrobial resistance mutations into 12 *E. coli* isolates using oligonucleotide-mediated mutagenesis [37]. The total number of mutant constructs was 67, after accounting for failed constructions and isolates that already harbored the mutation (Fig 1). The genetic backgrounds sampled were the laboratory strain MG1655 and 11 clinical isolates, including enterohemorrhagic (EHEC) and extraintestinal pathogenic (ExPEC) *E. coli*, that vary in serotype, antibiotic resistance profile, and plasmid presence [38,39]. The AMR mutations confer fluoroquinolone, rifampicin, or aminoglycoside resistance by modifying drug binding sites or by upregulating antibiotic efflux [13,40–42] and cause clinical resistance for pathogens such as *E. coli*, *Pseudomonas aeruginosa*, and *Mycobacterium tuberculosis* [12,26,43–47]. Thus, the mutant library includes a range of AMR mutations causing resistance to multiple antibiotic types introduced in genetic backgrounds that sample the genomic diversity of pathogenic *E. coli* populations found in nature.

**Fig 1.**
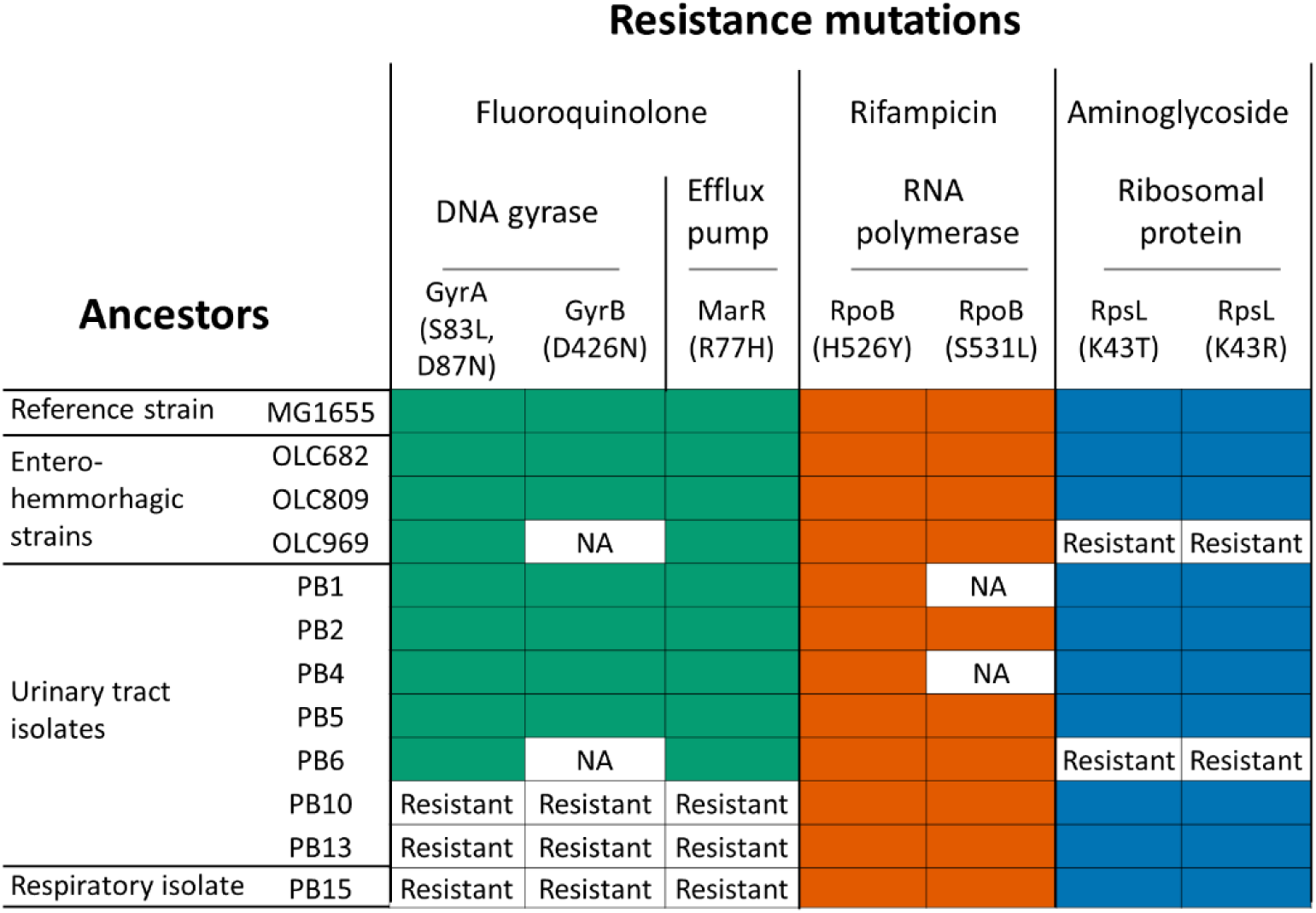
Library of *E. coli* isolates with introduced AMR mutations. The gene and specific amino acid change of the 7 introduced mutations are indicated. The mutations confer resistance to three antibiotic classes and alter the indicated cellular targets. The 12 *E. coli* genetic backgrounds include a common laboratory strain (MG1655) and 11 clinical isolates collected from patients in Canadian hospitals. 67 of the 84 potential mutation-by-genotype combinations were successfully constructed. 13 combinations were not attempted due to high intrinsic resistance of the ancestor (Resistant), and four combinations were unsuccessfully introduced (NA).

### Mutations cause similar increases in antibiotic resistance across *E. coli* genetic backgrounds

We expected the mutants to exhibit increased resistance to antibiotics whose inhibitory action or efflux was directly impacted by the introduced mutation. However, indirect effects of the mutations against non-target antibiotics (ie, cross-resistance or collateral sensitivity), and the extent to which antibiotic susceptibilities depended on genetic background were unknown. We therefore performed minimum inhibitory concentration (MIC) assays to quantify fold-changes in susceptibility to both target and non-target antibiotics (Fig 2). We found that the introduced mutations significantly increased resistance to target antibiotics across the genetic backgrounds. Thus, fluoroquinolone resistance mutations (in *gyrA*, *gyrB*, and *marR*) increased ciprofloxacin resistance, *rpoB* mutations increased rifampicin resistance, and *rpsL* mutations increased streptomycin resistance. Although several instances of collateral sensitivity were observed, none were significant when considering all genetic backgrounds.

**Fig 2.**
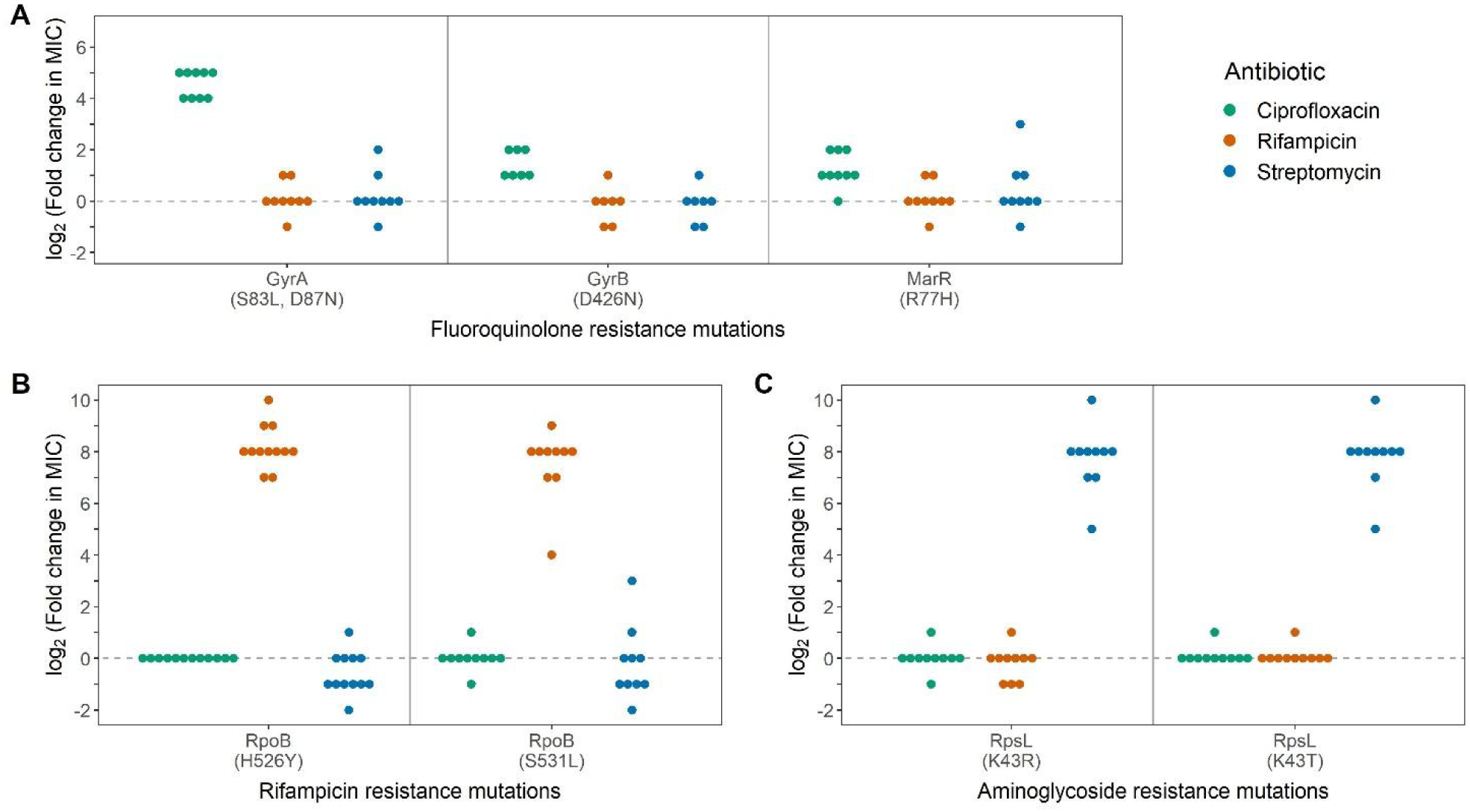
Mutations introduced in different genetic backgrounds consistently increase resistance to target antibiotics. Median fold-changes in antibiotic susceptibility [log_2_ (MIC_mutant_ / MIC_ancestor_)] are plotted for mutants with introduced (A) fluoroquinolone, (B) rifampicin, and (C) aminoglycoside resistance mutations. MICs were determined for three antibiotics (one target and two non-target) in triplicate. Individual points within each category represent different genetic backgrounds with the same introduced mutation. Each of the mutations significantly increased resistance to target antibiotics across genetic backgrounds (multiple comparison t-test with Bonferroni correction; p_adj_<0.05) with no significant effects on susceptibility to off-target antibiotics. The collateral streptomycin sensitivity observed for the RpoB (H526Y) mutants was not significant when adjusting for multiple comparisons (p_adj_=0.54).

There was little variation in the fold increases in antibiotic resistance for mutants sharing the same mutation but differing in genetic background. In a mixed effect linear model, knowledge of the identity of the mutation explained 89% of the fold change variance, with random effects of genetic background contributing only 9.7% of the explained variance. Moreover, some of the variation in MIC fold increase to target antibiotics could be explained by differences in the initial resistance of the ancestral isolates. For one mutation (RpoB (H526Y)), there was a negative correlation between fold increase in rifampicin MIC and the initial resistance of the ancestor (S1 Fig). This result is suggestive of diminishing returns epistasis, where mutations confer smaller beneficial effects in more fit genotypes [48], although the generally low MIC variation among the isolates prevented a robust test of this phenomenon. In conclusion, analysis of the antibiotic susceptibilities found that the introduced AMR mutations predictably increased AMR to target antibiotics across genetic backgrounds with few collateral effects against non-target antibiotics.

### Fitness effects of mutations vary widely across genetic backgrounds and growth environments

We next investigated whether the consistent increases in antibiotic resistance caused by the mutations were also reflected in predictable competitive fitness effects in antibiotic-free environments. Though AMR mutations are generally expected to be costly, fitness effects could vary depending on the identity of the mutation, genetic background, or growth environment. We measured fitness effects in head-to-head competition assays in four discrete antibiotic-free growth environments: rich (LB) and minimal (M9-Glucose) laboratory media and two media that simulate urinary tract and colon environments colonized by pathogenic *E. coli* [49,50]. The growth yield of the environments varied, with an over 10-fold change in carrying capacity between the highest yield (LB) and lowest yield (Urine) environments (S2 Fig). Fitness effects were estimated by calculating the fitness of each mutant relative to its unmutated ancestor, thus allowing for comparisons between mutants generated from different genetic backgrounds.

The fitness effects of the AMR mutations are summarized in Fig 3, where individual points in each boxplot represent different genetic backgrounds sharing the same mutation. The mutations were generally costly (with relative fitness < 1); however, there was wide variation in the fitness effects including neutral and, surprisingly, even beneficial effects. In contrast to the antibiotic susceptibility phenotypes, the mutations frequently exhibited large variation in fitness effects across genetic backgrounds. The dependence of the fitness effects on genetic background is evidence of epistasis between AMR mutations and genetic backgrounds, a type of G X G interaction.

**Fig 3.**
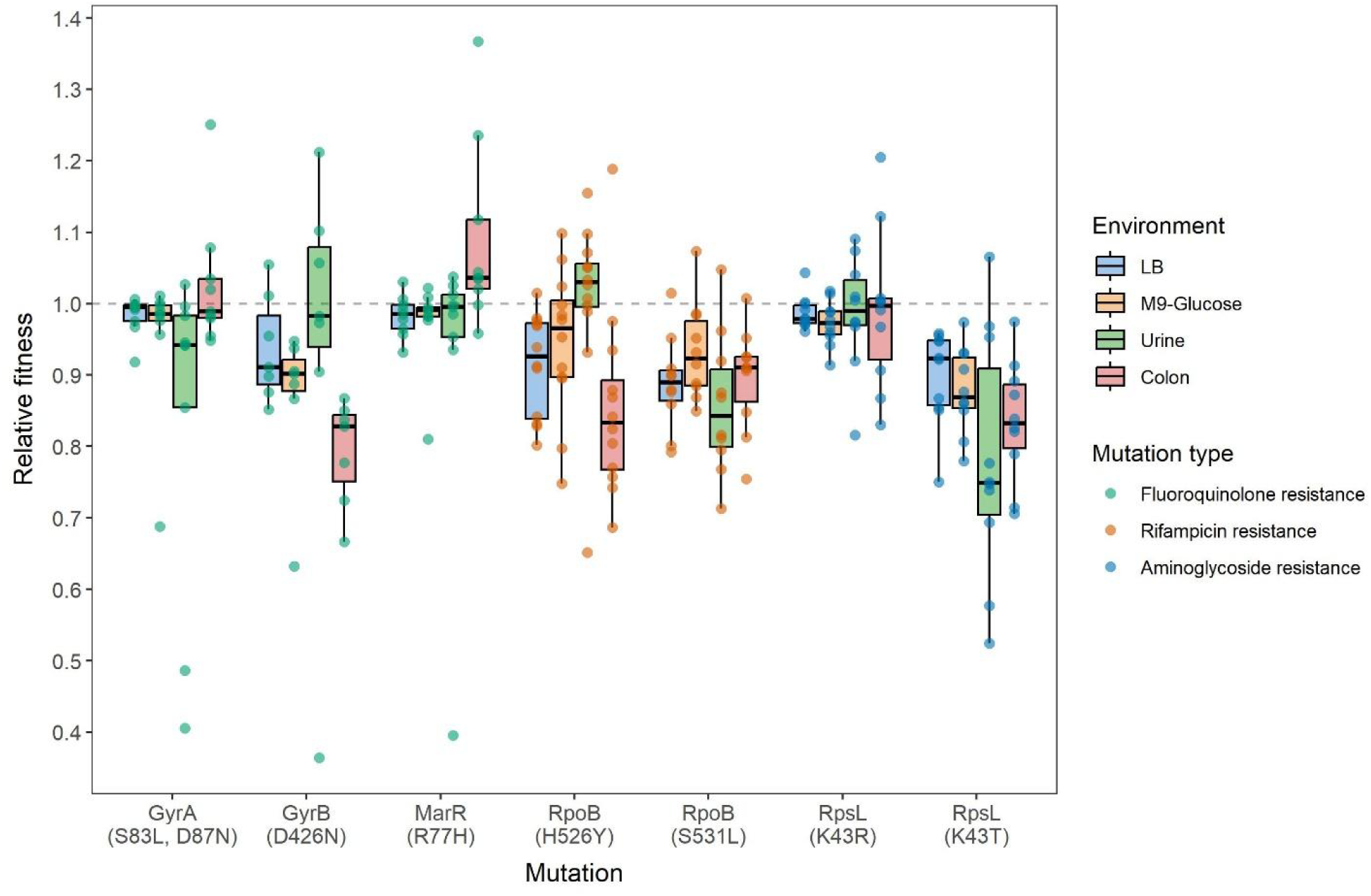
AMR mutations exhibit wide variation in fitness effects across genetic backgrounds and environments. The fitness effects of AMR mutations were determined in four antibiotic-free environments: LB, M9-Glucose, synthetic urine, and synthetic colon media. The data are grouped by mutation and environment, with individual points indicating fitness effects measured in different genetic backgrounds. The boxplots summarize the distributions of fitness effects (median, first and third quartiles, and nonoutlier minimum and maximum values). Additional fitness effect information including genetic background identities, relative fitness values, and significance levels is provided in S3 Fig.

We summarized the fitness effects for all the mutation and genetic background combinations in each environment with three statistics: (1) the mean fitness effects across mutations and genetic backgrounds; (2) the overall variance in fitness effects; and (3) the amount of epistasis between mutations and genetic backgrounds (Fig 4). Epistasis was estimated using the summary statistic gamma (γ), defined as the correlation of fitness effects of the set of AMR mutations across multiple genetic backgrounds [51]. We found that despite similar mean fitness effects (between -0.06 and -0.092), the variance of the fitness effects and amount of epistasis varied considerably depending on the growth environment. Fitness effect variance was much larger in the non-standard environments (synthetic urine and colon media), whereas epistasis was stronger (i.e., there was a low fitness effect correlation) for synthetic urine and M9-Glucose media, compared to synthetic colon and LB media. Despite differences in the total strength of epistasis, the four environments exhibited roughly similar proportions of epistasis types (S4 Fig), with ∼60-70% classified as magnitude epistasis, in which genetic background affects fitness non-additively in the same direction, and ∼30-40% as sign epistasis, in which genetic background affects the direction of the fitness effect (e.g., deleterious to beneficial or vice versa). Taken together, our results demonstrate the important role of epistasis in determining fitness effects of AMR mutations and, furthermore, the influence of the growth environment on both the overall variation in fitness effects and strength of epistasis between mutations and genetic backgrounds.

**Fig 4.**
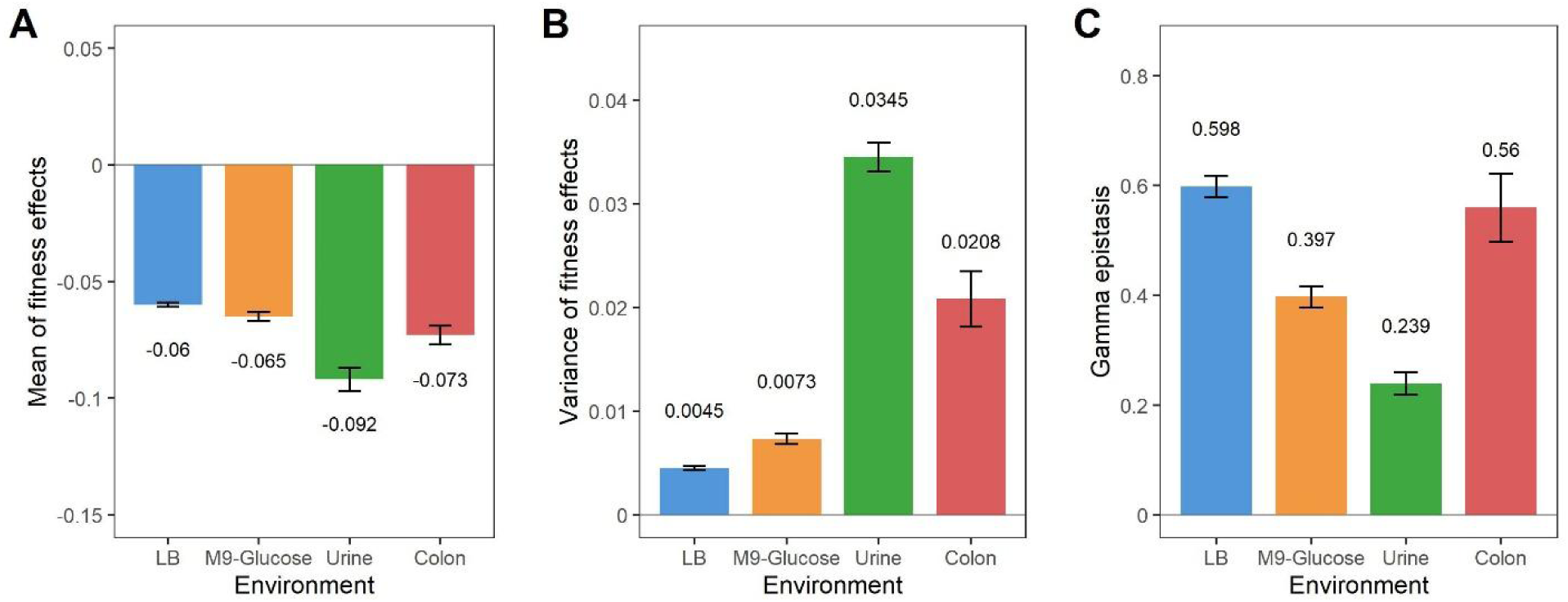
Fitness effects variance and epistasis between AMR mutations and genetic backgrounds differed between environments. The overall mean (A) and variance (B) of fitness effects of all mutation-genetic background combinations (83 mutants) were determined in each growth environment. Global epistasis (C) was estimated as gamma epistasis. Analysis of the amount and proportion of epistasis types (ie, magnitude vs. sign epistasis) underlying the epistasis measure are found in S4 Fig.

### Variation in fitness effects is governed by irreducible G x E interactions

The experimental fitness data highlight the dramatic influence of the growth environment on the fitness effects of AMR mutations. Changing the growth environment can lead to differences in fitness that depend on both the identity of the mutation and the genetic background (Fig 3). For example, the *gyrB* mutation was highly costly in synthetic colon media but less costly (and sometimes beneficial) in synthetic urine media. The *gyrA* and *marR* mutations, on the other hand, exhibited the opposite response in these two environments. These results demonstrate that the impact of growth environments on fitness can vary from mutation to mutation (ie, mutation by environment interaction).

The genetic background also played a major role in determining fitness effects across environments. Reaction norm plots of the fitness effects (S5 Fig) indicate that differences in a mutation’s fitness effect caused by shifting the growth environment depended on the genetic background, which impacts both the magnitude and direction (i.e., beneficial vs. deleterious) of the fitness effect. G x E interactions are also illustrated by the idiosyncratic genetic backgrounds that frequently yielded outlier fitness values depending on the introduced mutation and environment (ie, strains OLC682 and PB1). Overall, our results clearly illustrate that the fitness effects of the AMR mutations we sampled are influenced by complex interactions between AMR mutations, genetic backgrounds, and growth environment.

We next leveraged the factorial design of the experiment to quantify the importance of genetic background by environment (G x E) interactions in explaining the variation of fitness effects in the experimental data. We quantified the variance contributed by genetic background and environment in a linear mixed effect model treating genetic background, environment, and their interaction as random factors. We found that for each of the mutations, over 50% of the variation in fitness was explained by the interaction between genetic background and environment (Fig 5). Although the main effect of environment explained a portion (up to 17%) of the variation for several mutations, in general, the effect of environment on a mutation’s fitness effects strongly depended on its genetic background. In other words, neither knowledge of the growth environment nor of the identity of the genetic background were by themselves sufficient to predict fitness effects of each of the mutations.

**Fig 5.**
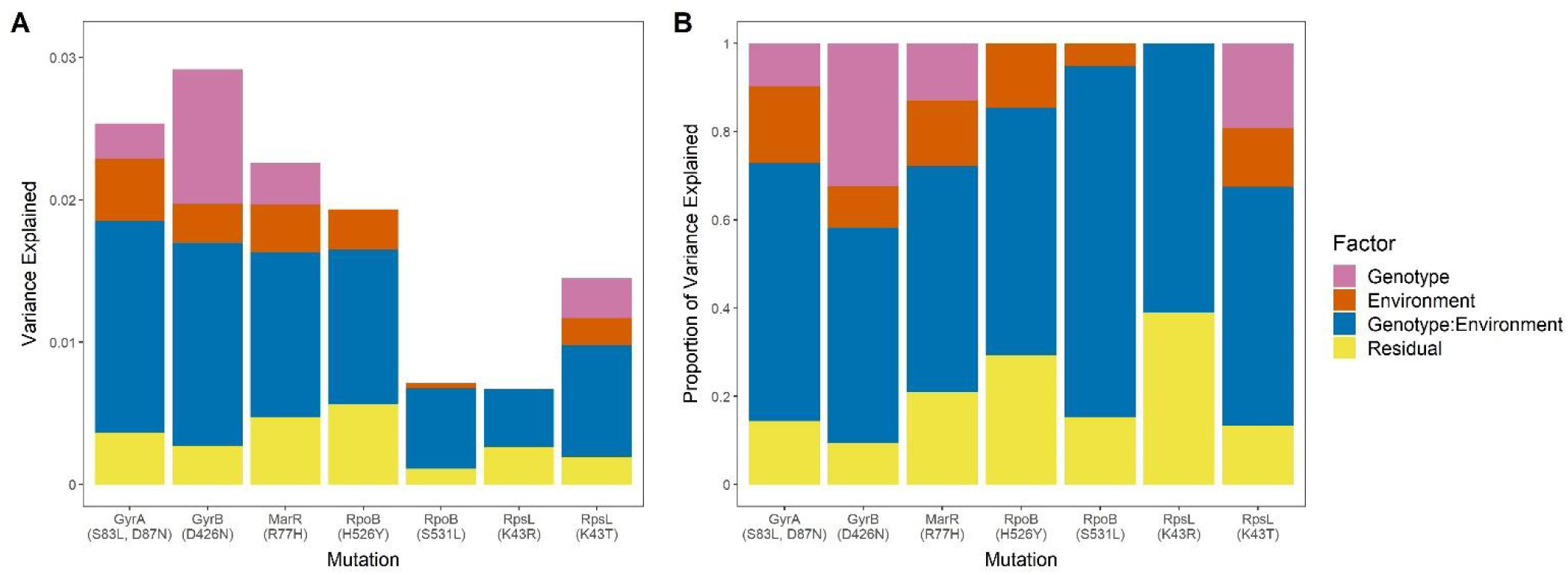
Genotype by environment interactions explain most of the variation in fitness effects for each AMR mutation. The experimental fitness effects data for each mutation was fit to a random effects model to determine the amount (A) and proportion (B) of variance explained by Genotype (ie, the genetic background), Environment, and the Genotype by Environment interaction. Reaction norm plots showing the responses of specific genotypes in each environment are found in S5 Fig. **AB**

### The complex G x G x E interactions in the experimental data are not captured by a probabilistic fitness landscape model

Our dataset provides a powerful test case for investigating the predictability of mutational fitness effects. For example, given data from one environment, can we predict the fitness effects of AMR mutations in a second environment? In the previous section we showed that the data exhibit strong variation in fitness effects and epistasis within and between environments, indicating ubiquitous G x G and G x E interactions (Figs 4 and 5). Would such variation be expected under a simple fitness landscape model, and would the data be consistent with the same fitness landscape being sampled independently for each of the environments? If yes, this indicates that at least statistical properties of the data (such as the variance in fitness effects and epistasis) are predictable, even in the absence of detailed mechanistic knowledge of the cellular and physiological effects. We chose the Rough-Mount-Fuji (RMF) model [34] to address this question due to its success in describing single environment fitness landscapes and its reliance on few parameters [36,52]. The model considers a genotype as a set of alleles at diallelic loci that each contribute additively to fitness plus a random epistatic component, specific to each genotype, that also contributes to fitness. The model can be tuned from completely additive to completely epistatic by adjusting the distributions from which the additive and epistatic components of fitness are drawn.

To test whether the experimental data were consistent with an underlying RMF landscape, we simulated a total of 100,000 fitness landscapes with 7 diallelic loci on 12 different genetic backgrounds (i.e., 84 genotypes) for 10,000 sets of model parameters (s_a and s_b), encompassing a total of 10^9^ simulated fitness landscapes (see Methods section for details). We then computed the three fitness statistics (Fig 4; mean fitness, fitness variance, and gamma epistasis) of the sampled data. We first computed which set of model parameters could best reproduce the fitness statistics observed in the experimental data for each environment separately, and how well this best model fits the data. Each statistic of the experimental data was generated under the RMF with the given parameters. Fig 6A shows a projection of the log-likelihood of each statistic, where the other dimensions are fixed for the parameters that provide the overall best fit. For LB and M9-Glucose environments, we found similar RMF model parameters as best fits for the experimental fitness statistics. The parameters that described Urine and Colon best were very different from the previous environments, requiring a much larger variance in the epistatic component of the model for both environments and also a larger variance in the additive component in the case of Urine. This discrepancy made it difficult to find shared model parameters to characterize the entire dataset.

**Fig 6.**
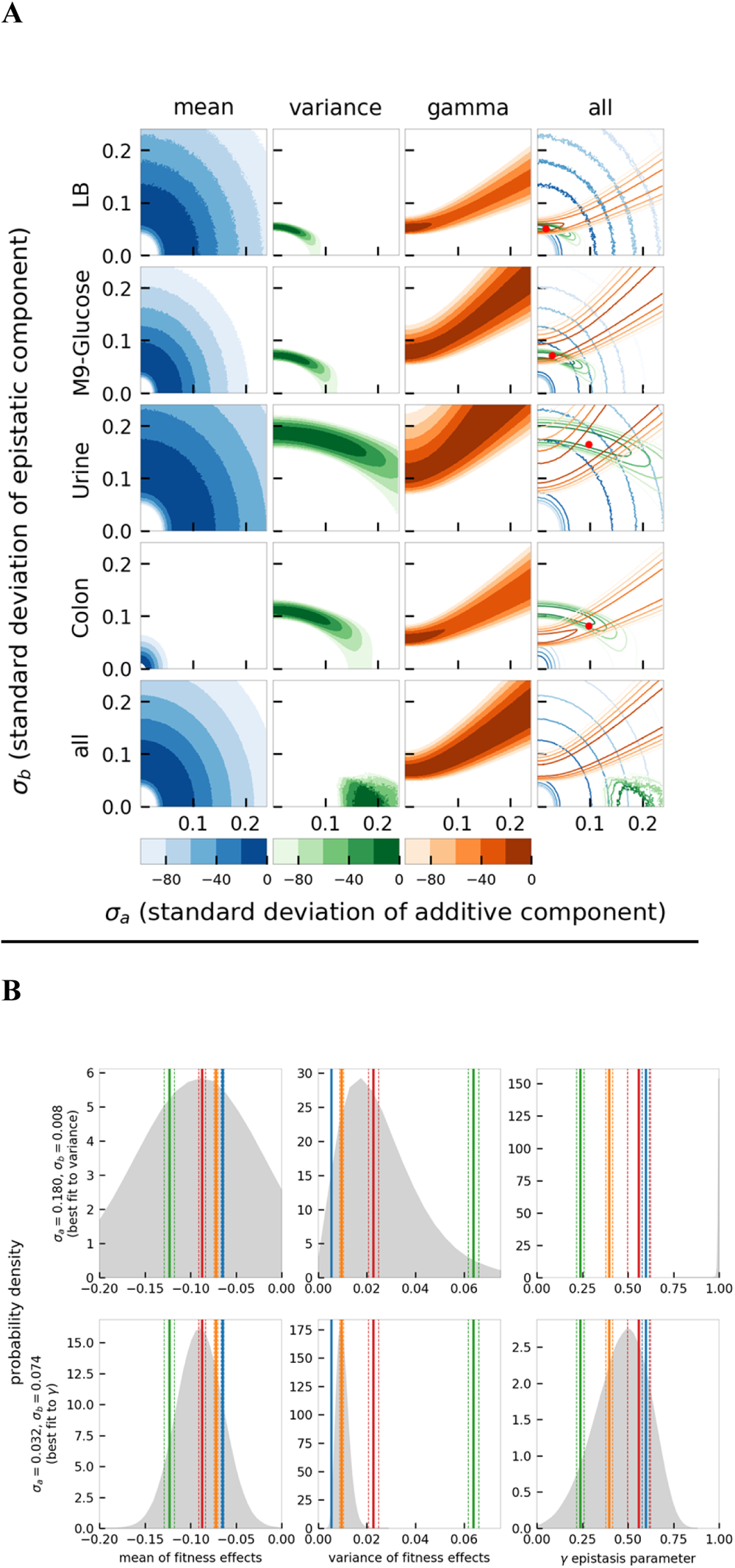
A Rough Mount Fuji genotype-fitness model only partially reproduces the experimental fitness effect statistics. (A) Log-likelihood surfaces for the mean of fitness effects, the variance of fitness effects, and the gamma epistasis parameter of the experimental data under a RMF model. x-axis represents the variance of the additive component (σ_a_) and y-axis the variance of the epistatic contribution (σ_b_). Each row shows log-likelihood surfaces for single environments and the last row for the conjugation of the four environments. The red dot represents an analytical estimate of the best parameters based on the mean values of the experimental data statistics only. The mean of the additive fitness effects (model parameter μ_a_) was fixed to the experimentally measured mean. (B) The RMF model cannot simultaneously capture the distribution of the variance of fitness effects and the epistasis for the four environments. The figure shows the distribution of the mean of fitness effects, the variance of fitness effects, and the gamma parameter under a RMF model. The top row represents a RMF landscape with parameters optimized to describe the variance of fitness effects and the bottom row optimized to describe the gamma parameter. The vertical colored lines represent the experimental value for each environment, with the dashed lines delimiting one standard deviation. The blue corresponds to the LB environment, the orange to M9-Glucose, the green to Urine, and red to Colon.

We next tested how well the model could capture the four different environments simultaneously. In other words, could the fitness effects of genotypes across different environments be explained by independent samples of the same underlying RMF landscape? Notably, the model failed entirely to accommodate the multi-environment data. Fig 6B illustrates the tradeoff that occurs when optimizing the parameter space for fitness variance vs epistasis by overlaying the model-generated fitness statistics (grey probability distributions) with the experimental values obtained for each environment (vertical lines). The results demonstrate that when optimized for fitness variance, the model completely fails to produce the epistasis values associated with any of the four environments in the experimental data. Conversely, optimizing the model for epistasis fails to capture the fitness variance. Thus, no single set of RMF parameters could adequately capture key features of the data across all environments.

### Fitness effects weakly correlate across pairs of environments

The complex G x G and G x E interactions observed in the experimental data, as well as the inability of the RMF model to accommodate the multi-environment dataset indicate that fitness effects measured in one environment do not necessarily translate to alternative environments. To directly ask whether fitness data from one environment can be used to predict fitness in another environment, we calculated the correlations of fitness effects between pairs of environments. Grouping all mutations together (Fig 7A), we found that, in general, fitness effects correlated poorly between environments (Pearson’s r < 0.2), with only one pair (M9-Glucose and Urine) showing a modest correlation.

**Fig 7.**
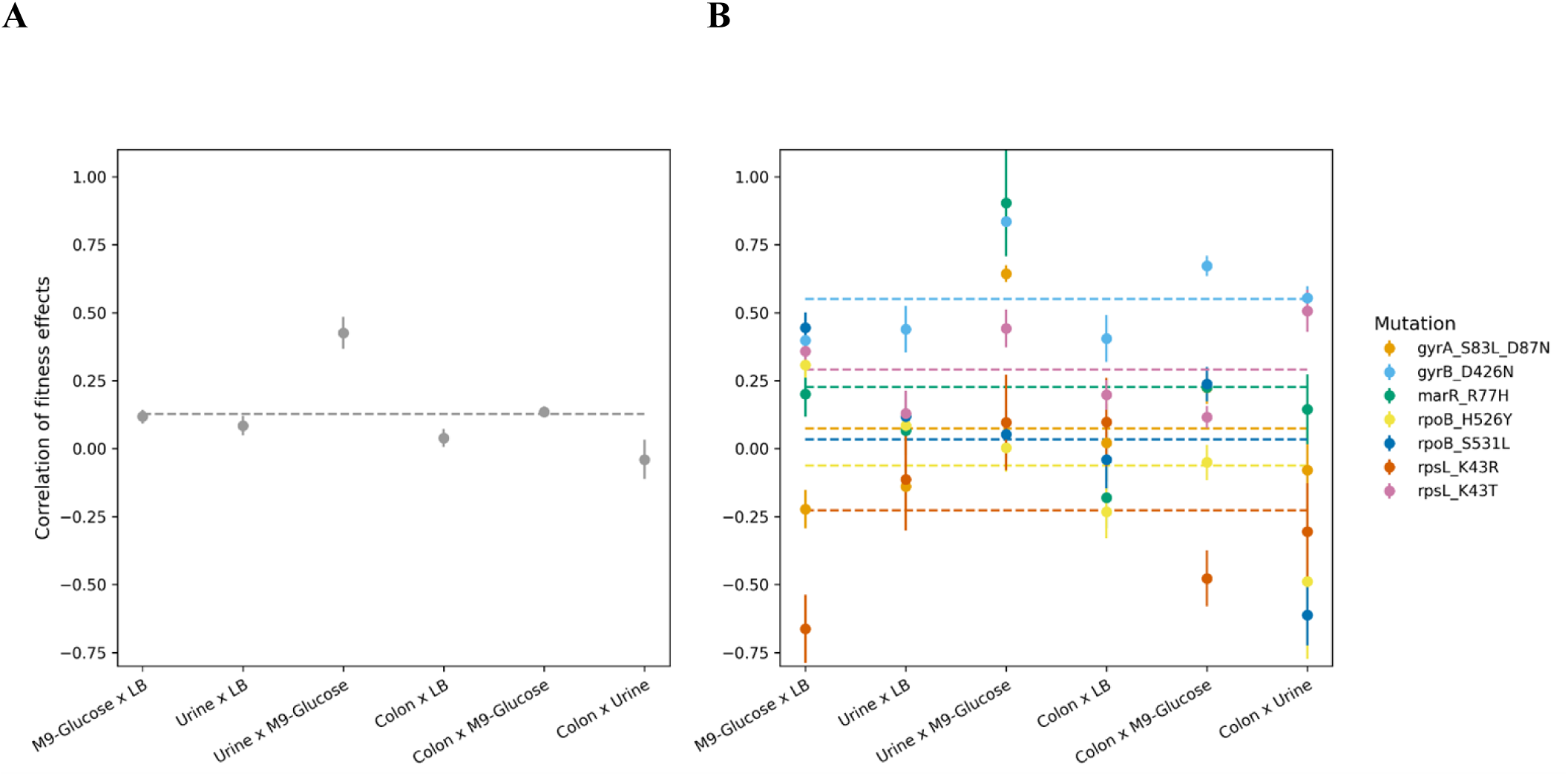
Fitness effects of the AMR mutations correlate weakly between pairs of environments. We determined the correlation of fitness effects of mutations when measured in different environments. For each environmental pair, the correlation coefficients (Pearson’s r) are plotted when including all mutants in the library (A), or when grouped by introduced mutation (B). Error bars indicate a one standard deviation range and horizontal lines indicate the mean of the correlation coefficients for all 6 comparisons. Correlation scatterplots, correlation coefficients, and significance values are provided in Figs S6 and S7.

Despite the poor overall correlations, it is possible that some individual mutations might show stronger relationships than others. Therefore, we disaggregated the data and determined the correlation of fitness effects for each mutation (Fig 7B; S6 Fig). Each data point in Fig 7B represents the correlation in fitness effects between two environments for an AMR mutation introduced into multiple genetic backgrounds. In general, the mutation-specific correlations were highly dependent on the environments under comparison, with only *gyrB* mutations yielding consistently positive correlations. The strongest overall environmental correlation, between M9-Glucose and synthetic urine, was driven by positive correlations for the fluoroquinolone mutations (*gyrA, gyrB,* and *marR* mutations). In contrast, the lack of correlation observed between synthetic urine and synthetic colon fitness effects was driven by a combination of positive and negative mutation-specific correlations, suggesting that genetic backgrounds associated with higher costs in synthetic urine media were associated with reduced costs in synthetic colon media and vice versa.

Variation in the strength of correlation between environments is indicative of mutation by genetic background by environment (G x G x E) interactions. Mutations with high correlations in fitness effects can be considered to have low levels of genetic background by environment (G x E) interaction, ie, different genetic backgrounds respond similarly to both environments. On the other hand, a low correlation indicates high levels of G x E interaction, ie, environmental effects on fitness depend on the genetic background. By this logic, the differences in correlations that we observe between mutations are indicators of G x G x E interactions, since the level of G x E interaction changes depending on the mutation. Although our sampling of genetic backgrounds is too sparse to make strong claims about mutation-level correlations, the variation in correlation coefficients that we observe in Fig 7B further supports that G x G x E interactions underlie the AMR mutation fitness effects.

### Genetic background fitness and phylogenetic relatedness are poor predictors of fitness effects

Although the experimental data show that genetic background is a significant source of variation on AMR fitness effects, further information about the isolates might help untangle the genetic background effects. We investigated whether two properties of the genetic backgrounds might explain the variation in fitness effects: their comparative relative fitness in each environment, and their phylogenetic relatedness. Beneficial mutations are expected to have smaller effect sizes for starting genotypes closer to a fitness peak (i.e., well-adapted to the growth medium) than genotypes further from the peak [53] . However, less is known about the expected magnitude of fitness effects for deleterious mutations at different distances from the peak [48]. Therefore, we investigated whether any correlation existed between the starting fitness of the genetic backgrounds and the fitness effects of the introduced AMR mutations. We estimated genetic background fitness in each of the four growth environments by competing each of the 12 unmutated isolates against a common competitor (S8A Fig). We observed some variation in the background fitness of the isolates, suggesting that some genotypes were better adapted to each growth environment than others. However, apart from a weak positive correlation in the M9-Glucose environment (Figs 8A and S9), we found no evidence that background fitness could predict the fitness effects of AMR mutations.

**Fig 8.**
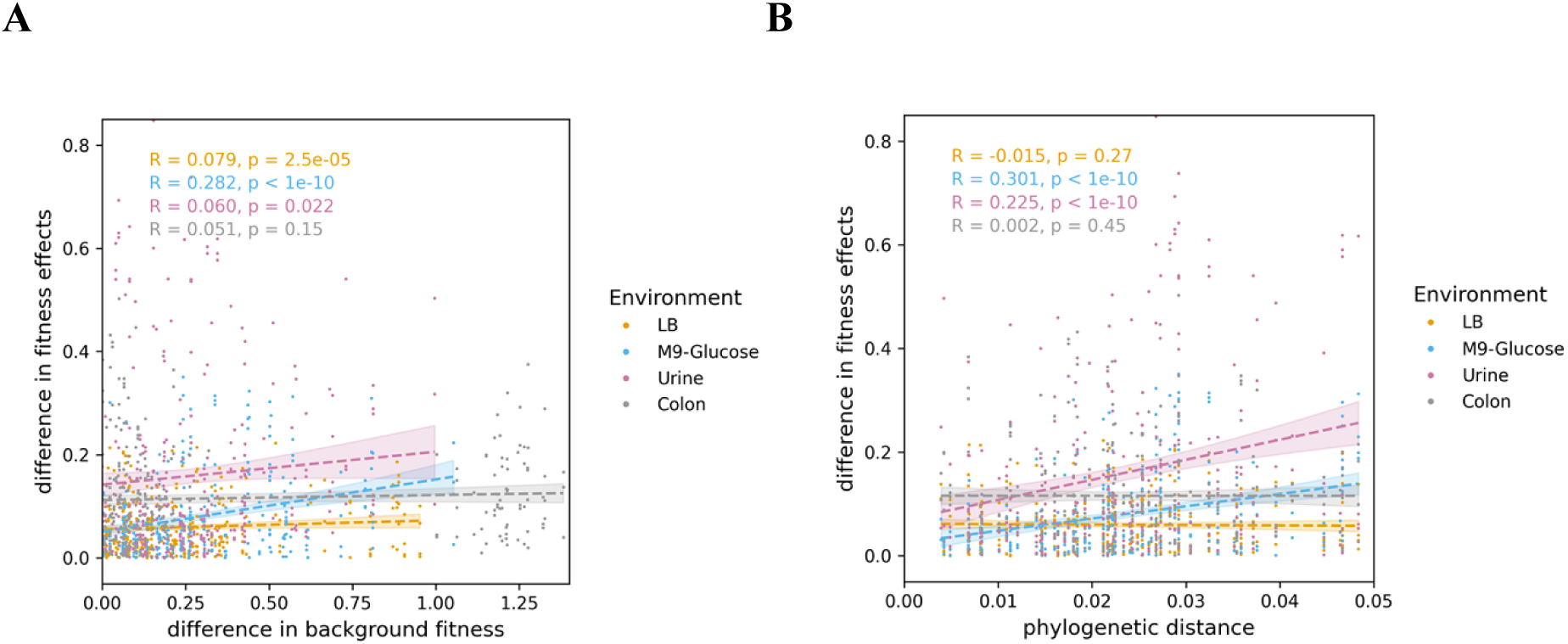
The fitness effects are not explained by differences in fitness or phylogenetic relatedness of the ancestral genetic backgrounds. The results of all pairwise comparisons between the 83 AMR mutants are shown, with difference in fitness effects plotted against difference in background fitness (A) or phylogenetic distance (B) for each comparison. Genetic background fitness was measured in head-to-head competitions between the twelve *E. coli* ancestors and a common competitor in each of the four growth environments (S8A Fig). Phylogenetic distance was estimated from a phylogeny based on whole genome alignments of the *E. coli* isolates (S8B Fig). The correlation lines and 95% confidence intervals are depicted for all mutants tested in each environment. Figs S9 and S10 show the correlations grouped by mutation.

We also tested whether phylogenetic relatedness could predict differences in fitness effects observed between genetic backgrounds. We reasoned that closely related genetic backgrounds would be more likely to share mutations that interact with the introduced AMR mutations, and would hence exhibit more similar fitness effects. We found weak positive correlations between differences in fitness effects and genetic distance for the M9-Glucose and Urine environments (Fig 8B), but no correlation for the LB and Colon environments. Interestingly, a subset of mutations was responsible for the positive correlations observed for the M9-Glucose and Urine environments (S10 Fig). Nevertheless, despite these exceptions, our analysis suggests that genetic background fitness and phylogenetic relatedness were poor overall predictors of AMR mutation fitness effect variation.

## Discussion

The predictability of evolutionary processes depends crucially on the impact of environmental and genetic variation on the fitness effects of mutations. Given the threat of antimicrobial resistance (AMR) to human health, knowledge of the factors determining the fitness effects of AMR mutations has important implications for antimicrobial stewardship. To the extent that resistance mutations are generally costly, antibiotic restriction is expected to reduce the prevalence of resistance. However, if costs of resistance are highly variable because they depend on environment and/or genetic background, then resistance might persist in favorable environmental or genetic refuges. Thus, we sought to measure, and ultimately predict, the effects of genetic background and environment on the fitness of AMR mutants.

We systematically measured the fitness effects of 7 resistance mutations across a range of *E. coli* genetic backgrounds and environments. The 12 genetic backgrounds included a standard laboratory strain and 11 clinical isolates, and the four growth environments included standard laboratory media, as well as media designed to mimic important sites of infection for *E. coli*. AMR mutations caused fairly uniform increases in resistance itself (Fig 2), and were on average costly in the absence of antibiotics (Fig 3). However, the magnitudes of their impacts on fitness were highly variable (Fig 3). Importantly, these fitness effects could not be predicted simply by knowing the identity of the resistance mutation, but instead depended to varying degrees on the assay environment, the genetic background of the host strain, and interactions between individual terms. These complex interactions rendered the fitness effects of resistance mutations highly unpredictable.

Genotype-by-environment interactions are well documented in the quantitative and evolutionary genetics literature. Across a broad range of organisms, mutations may have drastically different effects in different environments (reviewed in [54,55]). Likewise, there is growing evidence that AMR mutations, although typically deleterious, can have widely varying fitness effects depending on the environmental context [25,56–58]. We similarly find that the fitness effects of resistance mutations depend on the assay environment – a given mutation may be deleterious in some environments but neutral on average in others (Fig 3, GyrB (D426N)), or even beneficial in some environments but not others (e.g., the beneficial effect of MarR (R77H) in artificial colon). Although not the focus of the study, several genotype-by-environment interactions we observed have biologically plausible mechanisms based on the presence of specific constituents or nutritional complexity of the media. For example, *marR* mutations, which cause elevated expression of the AcrAB-TolC multidrug efflux pump [13,59,60], could be beneficial in synthetic colon media due to increased efflux of bile salts, a known substrate of the pump [61]. Similarly, differential fitness effects of RpoB (H526Y) mutations are thought to reflect alterations to global transcription that are beneficial specifically during growth in nutritionally poor media [62].

In addition to widespread G x E, we found that the fitness effects of AMR mutations depend on genetic context. In contrast to the relatively uniform effects of the mutations on resistance (Fig 2), the fitness effects in the absence of antibiotics could be deleterious, neutral, or beneficial depending on the genetic background. Our results reinforce the conclusions of empirical studies demonstrating the influence of genetic background on AMR fitness costs [24]. Trindade et al. [26] observed widespread epistasis among AMR mutations in *E. coli*, with combinations of resistance mutations introduced in the same genetic background frequently exhibiting non-additive fitness effects. Similarly, but at a larger phylogenetic scale, Vogwill et al. [23] identified large variation in the costs of rifampicin-resistance mutations across 8 different species within the genus *Pseudomonas*, with much of the variance attributed to the interaction between mutation and genetic background. Taken together, these experimental studies suggest that the costs of AMR are strongly influenced by epistatic interactions between AMR mutations and other loci over broad scales of relatedness, ranging from single nucleotide to strain and species level differences.

Our multi-factorial study design also allowed us to detect three-way interactions between mutation, background genotype, and environment (Fig 7). Here, different resistance mutations demonstrate contrasting G x E interactions. For example, fitness effects across genetic backgrounds are well correlated for MarR (R77H) between synthetic urine and M9-Glucose (ie, low G x E), but not for RpoB (H526Y) (ie, high G x E). Relatedly, we also find that overall levels of epistasis between mutations and genetic backgrounds differ from one environment to another (Fig 4C). Crucially, G x G x E interactions make it difficult to predict costs of resistance between environments, implying that mutation-genotype combinations will respond idiosyncratically to a change in environment. Thus, estimates of fitness using standard lab strains or growth environments may be poor predictors of fitness in clinical settings, limiting our ability to predict which types of resistance will respond to antibiotic restriction, and how quickly they will do so.

The complex gene by environment interactions underpinning the mutational fitness effects in our experimental dataset provided a test case for the ability of a fitness landscape model to reproduce the observed patterns in the data. For this purpose, we chose one of the simplest probabilistic fitness landscape models, the Rough Mount Fuji (RMF) model, which features tunable epistasis with few parameters and thus lends itself as a base model to test hypotheses regarding the consequences of epistasis in evolution [35]. Although the RMF model does not incorporate any explicit expectations of how a fitness landscape may differ between environments, its probabilistic nature results in a large variation of fitness landscapes that can be produced under the same parameters. Therefore, we tested 1) how well the model could fit the data of each single environment, and 2) whether it could accommodate the data from all environments given the same set of parameters. We found that the Rough Mount Fuji model was able to fit the statistics of individual environments well (Fig 6), considering the variation in fitness effects observed across genetic backgrounds. The model parameters suggest substantial epistasis for all environments, with the standard deviation of the epistatic contribution to the fitness effects typically larger than the standard deviation of the mean additive contribution. Although RMF could accommodate the fitness effect statistics for each environment separately, we found that no set of parameters could successfully explain the same statistics when all environments were considered together. In other words, the fitness landscapes in different environments could not be described as independent draws from a model with common parameters. We conclude that because even the most general features of the underlying fitness landscape (i.e., the average additive and epistatic contributions to fitness) are irreconcilable between environments, successful prediction of fitness will require sophisticated models that incorporate additional, environment-specific factors.

The failure of our RMF-like model to predict fitness across environments raised the question of what additional information would be required. We investigated two possibilities here – phylogenetic relatedness, and relative fitness of the ancestral genotypes. Phylogenetic relatedness could in principle help to predict epistatic interactions, since closely related genotypes share more polymorphisms than distant relatives. The fitness effects of focal resistance mutations should be similar for close relatives to the extent that these effects are modulated by these shared polymorphisms. However, we found no correlation between relatedness and the fitness effects of resistance mutations (Fig 8B). One potential explanation for this lack of correlation is that mutations that interact with our focal resistance mutations arise frequently, and are thus not shared even between closely related genotypes.

Alternatively, the fitness effects of a mutation may not be determined by specific interactions with other mutations in a given genetic background, but rather by global properties of a genotype. Fitness is a clear candidate for such a property – in the widely observed phenomenon of diminishing returns epistasis, beneficial mutations confer a smaller gain for genotypes with higher starting fitness [53,63–65]. Wang et al. [53], for example, provided clear evidence that the fitness effects of several beneficial mutations were predicted well by the fitness of the ancestral genotype, but not by relatedness or by metabolic similarity. We found some evidence for diminishing returns epistasis for level of resistance (S1 Fig) – that is, for the trait towards which these mutations provide a direct benefit. However, we found no relationship between ancestral fitness and mutational effects on fitness in the absence of drug (Fig 8A), such that indirect effects did not show a pattern of diminishing returns. Indeed, while there is theoretical justification of diminishing returns epistasis for beneficial mutations from Fisher’s geometric model (e.g., [66]), expectations are unclear for non-beneficial mutations.

We suggest that predictability may be improved by specific information concerning the assay environments. Characteristics of environments that could be quantitatively compared, such as nutrient concentrations, could be valuable to inform the direction in which the model should be expanded to account for differences between environments. It is worth noting that, overall, fitness effects were best correlated between the M9-Glucose and synthetic urine environments (Fig 7A). These environments offer lower nutritional complexity than do LB and synthetic colon media, both of which contain relatively large amounts of complex nutrient mixtures (e.g., tryptone). Further exploration of the impact of nutritional environment on predictability may thus be warranted.

In conclusion, this study provides a systematic view of the impact of both genetic background (i.e., epistasis) and environment (i.e., G x E interactions) on the fitness effects of AMR mutations. In the context of antimicrobial stewardship, our results suggest that the response of resistant bacteria to antibiotic restriction might be difficult to predict. The outcome of a restriction protocol might depend on the genetic background(s) of the resistant microbial population and on the availability of environment refuges where costs are diminished. Nevertheless, although we found that in some genetic and environmental contexts AMR mutations were neutral or beneficial, overall the mutations tended to be costly and variation in fitness effects was driven more by differences in the magnitudes of the costs rather than changes in sign from costly to beneficial (S4 Fig). Thus, to the extent that our findings translate to clinical settings, we would expect antibiotic restriction interventions to be successful on average in reducing the prevalence of resistance, but at an unpredictable pace. Furthermore, some mutations were more consistently costly across environments and genetic backgrounds (i.e., RpsL (K43R) and RpoB (S531L)), suggesting that knowledge of the mutation could provide some level of predictive value for antibiotic restriction outcomes. In addition, our study calls for the further development of fitness landscape models across environments and their evaluation in the light of data such as those presented in this study. Such models could help identify the variables that influence predictability and inform subsequent experimental study design.

## Materials and Methods

### Bacterial strains, growth conditions, and antibiotics

The *E. coli* isolates sampled for AMR mutagenesis include the K-12 reference strain (MG1655) [67], six extra-intestinal isolates collected from patients during the 2007-11 CANWARD survey of antibiotic-resistant pathogens in Canada [38,68], and three enterohemorrhagic strains from the Ottawa Laboratory Carling culture collection [39]. Additional *E. coli* strains used for molecular cloning procedures include DH5α λpir and WM3064 [69].

Bacteria were routinely cultured at 37 °C in LB broth [70] with shaking (150 rpm) and plated on LB with 1.5% agar. Antibiotics were added to LB media from stocks prepared at the following concentrations: Ampicillin (100 mg/ml in water), Ciprofloxacin (10 mg/ml in 0.1 M NaOH), Rifampicin (50 mg/ml in DMSO), Streptomycin (100 mg/ml in water). Media were also supplemented when appropriate with 0.2% Arabinose (from sterile-filtered 10% solution in water), 5-10% Sucrose (from sterile-filtered 50% solution in water), 0.3 mM Diaminopimelic acid (DAP; from 60 mM solution in water), 1 mM IPTG (from 100 mM solution in water), 40 µg/ml X-Gal (from 20 mg/ml solution in Dimethylformamide).

### Construction of AMR mutants by site-directed mutagenesis

AMR mutations were introduced into *E. coli* genomes by oligonucleotide-mediated recombineering as described by Lennen et al. [37]. The Lambda Red plasmid (pMA7-SacB) encodes arabinose inducible Lambda Beta and Dam methylase functions. Lambda Beta promotes chromosomal recombination of single-stranded DNA with short (< 100 base pair) regions of homology [71], while Dam methylase induction increases recombination efficiency by transiently disabling the *E. coli* DNA mismatch repair system [37]. Recombineering oligonucleotides were designed with the MAGE Oligonucleotide Design Tool (MODEST) [72], and encoded AMR point mutations centered within 90 bp of sequence homologous to *E. coli* (MG1655) genomic sites (S1 Table). Oligonucleotides were synthesized by Integrated DNA Technologies (standard desalting with no additional purification) and suspended in water to 100 µM. pMA7-SacB was a gift from Morten Sommer (Addgene plasmid # 79968; http://n2t.net/addgene:79968; RRID: Addgene_79968). pMA7-SacB was transformed into *E. coli* strains by electroporation with selection on LB agar with 100 µg/ml ampicillin (or 6400 µg/ml ampicillin for the β-lactam resistant isolate PB6).

Site-directed mutagenesis was performed according to the protocol described by Lennen et al. [37]. Strains carrying pMA7-SacB were inoculated in LB broth with ampicillin, incubated overnight at 37 °C with shaking, subcultured (1:100) in LB with ampicillin, and incubated for another 2-3 h. Arabinose (0.2%) was added to the cultures to induce Lambda Beta and Dam methylase functions. Cultures were incubated for 15 min at 37 °C with shaking, then transferred to an ice bath and chilled for 15 min. Electrocompetent cells were prepared by pelleting 10 ml of culture, resuspending and pelleting the cells three times with ice-cold sterile water (twice with 5 ml and once with 1 ml), with a final resuspension in 0.25 ml ice-cold water. For each electroporation, 25 µl of competent cells were mixed with 25 µl of oligonucleotide (diluted to 4 µM in water) in an Eppendorf tube, transferred to a chilled 0.2 cm cuvette (Bio-Rad), and electroporated at 2.5 kV (Bio-Rad MicroPulser; Ec2 setting). Cells were suspended in 3 ml LB broth and incubated with shaking overnight in test tubes.

Mutants were selected by plating serial dilutions of the cultures on LB agar supplemented with the antibiotic corresponding to the introduced mutation (ciprofloxacin: 0.025-0.05 µg/ml; rifampicin: 25-50 µg/ml; streptomycin: 25-50 µg/ml). Successful transformations yielded higher colony counts (typically 10^4^ to 10^6^ cfu/ml) on the selective media in comparison to negative controls (i.e., cells electroporated with no added oligonucleotide). Candidate mutants were streaked on LB containing 5% sucrose to cure strains of the Red plasmid by SacB-mediated counterselection [37,73]. Successfully introduced mutations were identified by sequencing (Genome Quebec) PCR products amplifying the genomic locus targeted by the recombineering oligonucleotide (S2 Table).

The final mutant library comprised 67 sequence-confirmed mutants. Thirteen mutants were not constructed in cases where the ancestor already carried the mutation (*gyrA* mutations in PB10, PB13, and PB15) or had high intrinsic resistance that compromised the selection of recombinants (*gyrB* and *marR* mutations in PB10, PB13, and PB15; *rpsL* mutations in PB6 and OLC969). Four mutants failed to be generated despite their attempted construction (*gyrB* mutations in PB6 and OLC969; *rpoB* S531L in PB1 and PB4).

### Construction of YFP-marked *E. coli* strains

We marked *E. coli* strains with yellow fluorescent protein (YFP) to distinguish bacteria by fluorescence in competitive fitness assays. A constitutive YFP expression cassette was inserted in the chromosome of *E. coli* isolates, replacing *lac* operon sequences [74] spanning 62 bp upstream from the *lacI* start codon to 56 bp upstream from the *lacA* stop codon. We constructed a custom allelic replacement (AR) plasmid (pR6KT-SacB-Δ*lacIZYA*::YFP) with YFP sequences flanked by *lac* operon-targeting sequences on a suicide plasmid (pR6KT-SacB) [75,76] that replicates only in hosts carrying the lambda pir gene [77]. Selectable (ampicillin and tetracycline resistance genes) and counterselectable (*sacB*) markers on the plasmid facilitate two-step allelic exchange between plasmid and chromosomal sequences [19,76]. We used the primers listed in S2 Table to PCR-amplify the YFP cassette from a source plasmid (pAH1T-P*_A1/04/03_*-YFP) [75] and the *lac* targeting sequences from MG1655 chromosomal DNA. The three PCR products were spin-column purified and ligated to pR6KT-SacB in one-pot Golden Gate assembly reactions containing the Type IIS restriction enzyme BsaI and T4 DNA ligase, as described by Hinz et al. [75]. Assembly reactions were transformed into chemically competent *E. coli* DH5α λpir by the Inoue method [70], and ampicillin-resistant clones carrying the assembled YFP AR plasmid were identified based on PCR screening and detection of fluorescence in liquid cultures.

The allelic replacement procedure involved plasmid conjugation from donor to recipient *E. coli* strains, followed by two sequential *recA*-mediated recombination events between homologous sequences shared by the plasmid and recipient chromosome. The YFP AR plasmid was transformed into the auxotrophic donor strain WM3064, a pir-expressing *E. coli* that requires diaminopimelic acid (DAP) supplementation in growth media [69]. Conjugations were performed between the plasmid-bearing donor (grown in LB with 0.3 mM DAP and 100 µg/ml ampicillin) and recipient strains on LB agar containing 0.3 mM DAP. Exconjugates harboring chromosomally integrated plasmid were selected on LB agar containing 100 µg/ml ampicillin or 10 µg/ml tetracycline (selecting for recombinant recipients) and lacking DAP (preventing growth of the WM3064 donor). Selection for the second recombination event (loss of AR plasmid sequences) was accomplished by plating the exconjugates on LB agar containing 10% sucrose, to eliminate genotypes carrying the plasmid-borne *sacB* gene. Sucrose resistance could result from chromosomal excision of plasmid backbone sequences, leading to either replacement of the *lac* operon with the YFP cassette or reversion to the wild-type *lac+* sequence. These two possibilities were distinguished by including 1 mM IPTG and 40 µg/ml X-Gal in the sucrose selection media, enabling blue-white screening of *lac* genotypes. White (*lac-*) sucrose-resistant colonies were subsequently screened for loss of plasmid antibiotic resistance markers and presence of fluorescent signal.

The YFP cassette was successfully transferred into 7 *E. coli* genetic backgrounds: MG1655, PB1, PB4, PB13, PB15, OLC682, and OLC969. Introduction of the marker failed in the remaining 5 genetic backgrounds (PB2, PB5, PB6, PB10, OLC809) due to the inability to select for successful recombinants during the allelic replacement procedure.

### Minimum inhibitory concentration assays

Antibiotic susceptibilities of AMR mutants and their ancestors were determined by broth dilution Minimum Inhibitory Concentration (MIC) assays [78]. Two-fold serial dilutions of antibiotics were prepared in 100 µl of LB broth in 96-well plates, and 100 µl of *E. coli* overnight cultures (diluted 1:1000 in LB) were added to each well. Plates were incubated statically at 37 °C for 20 h, and the OD600 of each well was measured with a microplate reader (Biotek Synergy H1). MICs were defined as the lowest antibiotic concentration yielding an OD600 signal of 0.15 after subtraction of values for uninoculated blanks. The following antibiotics and two-fold concentration ranges were tested: ciprofloxacin (0.003125 to 204.8 µg/ml), rifampicin (3.125 to 3200 µg/ml), and streptomycin (3.125 to 3200 µg/ml). MICs for each of the antibiotics were determined for all 67 AMR mutants and their 12 ancestors in three replicate assays inoculated from the same overnight cultures. Two *rpoB* mutants [PB4 RpoB (H526Y) and PB15 RpoB (S531L)] and all except one *rpsL* mutant [PB13 RpsL (K43R)] grew at the maximum tested concentrations (3200 µg/ml) of rifampicin and streptomycin, respectively. Their MICs (6400 µg/ml) should be considered lower-bound estimates, and therefore, variation in streptomycin resistance among *rpsL* mutants was not accurately determined. Plots show the median fold change in MIC due to mutation for the three replicate estimates (calculated by dividing the MIC of the mutant by the median MIC of the ancestor).

Differences in antibiotic susceptibility caused by the introduced mutations across different genetic backgrounds were determined by a multiple comparison t-test in R Version 4.2.2 [79] with the compare_means function in the ggpubr package. Median fold-changes in MIC (log_2_-transformed) were compared between mutants and ancestors (with antibiotic as a grouping variable) and significance was assessed using Bonferroni-adjusted p-values. The predictability of MIC fold increases was determined by fitting a mixed-effect linear model using the lmer function in the lme4 package. The model predicts the fold-change in MIC (log_2_-transformed) for target antibiotics from the identity of the introduced mutation (fixed effect), with the genetic background and interaction between genetic background and mutation included as random effects. Correlations between MIC fold-increases and ancestral MIC were determined using stat_cor in the ggpubr package.

### Growth media and carrying capacity estimates

Competition experiments were performed in lysogeny broth (LB), M9-Glucose, synthetic urine media, and synthetic colon media. LB and M9-Glucose were prepared as described [70]. Synthetic urine media is a defined media containing 416 mM urea and 10 mM creatinine and was modified from published recipes [49,80] by the addition of Casamino acids to augment bacterial growth. Synthetic colon media is a tryptone-based medium supplemented with 0.4% Bile Salts and was prepared as described [50]. The concentrations of each media component and details on preparation are found in Figs S3, S4, S5, and S6.

The carrying capacity was assessed for each ancestral isolate following 20 h of growth in each competition growth medium. Isolates were inoculated in duplicate in LB broth and incubated 24 h at 37 °C with shaking. Cultures were diluted (1:100) into each of the competition growth media (LB, M9-Glucose, Synthetic urine media, and Synthetic colon media) and incubated 24 h for media acclimation. Cultures were diluted (1:100) into fresh competition growth media and incubated for 24 h. Growth yields were determined by plating serial dilutions of the cultures on LB agar, and calculating the number of colony forming units per ml.

### Competitive fitness assays

Relative fitness was determined in head-to-head competitions [12] between unmarked AMR mutant constructs and YFP-marked competitors co cultured in each growth environment (LB, M9-Glucose, Synthetic urine media, and Synthetic colon media). Focal strains were competed against isogenic YFP-marked competitors or, when the isogenic competitor could not be constructed, a common competitor (MG1655-YFP). Four replicate competitions derived from independently inoculated cultures were performed for each genotype by environment combination. Pure cultures of unmarked mutants, unmarked ancestral isolates, and YFP-marked competitors were prepared in 24-well microplates by inoculating 1.5 ml of competition growth medium with colonies from freshly streaked LB plates. After 24 h of incubation at 37 °C with 150 rpm shaking, 100 µl of each unmarked strain was mixed with 100 µl of its corresponding YFP-marked competitor. 15 µl of each mix was diluted in 1.5 ml growth medium (1:100 dilution), and the remainder was reserved for immediate flow cytometry analysis (initial timepoint). After 24 h of incubation at 37 °C, competition cultures were analyzed by flow cytometry (final timepoint).

The relative frequencies of unmarked and YFP-marked cells in initial and final timepoints of the competitions were determined by flow cytometry analysis. Cultures were diluted in 1 ml of freshly filtered (0.2 µm pore size) 1X M9 salts prior to analysis (1:1000 for LB, 1:500 for M9-Glucose, 1:50 for synthetic urine media, and 1:500 for synthetic colon media). Cells were counted using a Gallios Flow Cytometer (Beckman Coulter) with a minimum of 20,000 counts per sample. Fluorescence was detected using the 488 nm excitation laser and 525/40 nm detection filter. Numbers of fluorescent and non-fluorescent cells were estimated using Kaluza analysis software. Signal thresholds for distinguishing fluorescent from non-fluorescent cells were established for each YFP-marked genotype by analyzing pure cultures of marked and unmarked strains. To distinguish cells from non-specific particles, gates were drawn in forward vs. side scatter plots that maximized the proportion of YFP-positive counts in pure cultures. The number of unmarked cells in each sample were estimated by subtracting the number of YFP-positive gated counts from the total number of gated counts.

### Relative fitness calculations

Relative fitness (⍵) was calculated as previously described [12] from the initial (*i*) and final (*f*) counts of the unmarked focal (*n*1) and marked competitor (*n*2) strains:

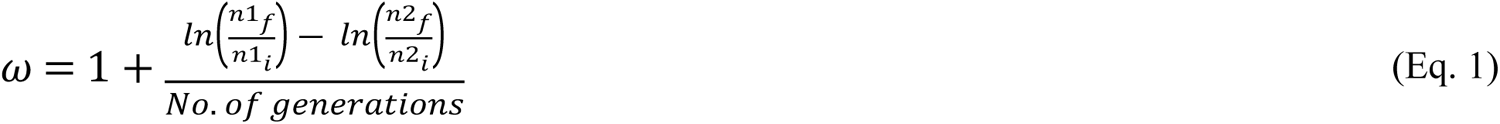

The number of generations was inferred from the 1:100 dilution factor, calculated as log_2_(100) ≅ 6.64. To account for fitness effects caused by the YFP marker or use of a non-isogenic competitor, the relative fitness value for each AMR mutant was divided by the relative fitness of its wild-type ancestor (competed against the same YFP-marked strain). These scaled fitness values, therefore, indicate the effects of the introduced AMR mutations alone.

Variance component analysis of the relative fitness estimates was performed in R using the lmer function in the lme4 package. For each mutation, a random effects model was fit that included genetic background, environment, and their interaction as random factors contributing to variance in relative fitness. The plots show the fitness effects variance explained by each random effect, as well as the proportion of total variance explained.

### Epistasis analysis

To quantify epistasis, we used the gamma statistic (γ), introduced by [51]. γ is defined as the average correlation of fitness effects across diverse genetic backgrounds in which a mutation manifests. This can be mathematically expressed as

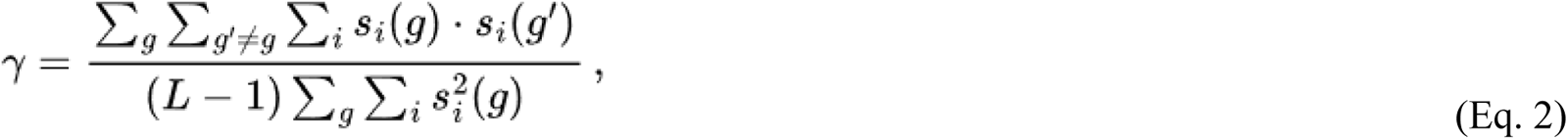

where *g* and *g’* index all possible genotypes, *i* all existing mutations, and *s_i_(g)* represents the effect of mutation *i* in the *g* background.

When the correlation is close to 1, the effect of the mutations is relatively consistent across the genetic backgrounds they appear, implying minimal epistasis. As the value of γ decreases, the correlation between mutation effects in different backgrounds weakens, indicating more pronounced epistasis. In the extreme, when the correlation approaches zero, the mutation effects in different genetic backgrounds become independent, signifying the highest level of epistasis.

In S4 Fig, we present an additional measure of epistasis, the difference in fitness effects when the genetic background is changed. We present its absolute value as this difference can be positive or negative. We further partition this epistasis into two categories: magnitude epistasis, where the direction of the fitness effect remains consistent, and sign epistasis, where a mutation transitions from advantageous to deleterious (or vice versa) when its genetic background is modified.

### Modeling

We employed a general version of the Rough Mount Fuji (RMF) model to try to capture the essential statistical features of the data. The model considers a genotype composed of *L* diallelic loci. Each allele is assigned an additive effect; on top of the additive effect, an epistatic component specific to each genotype is added. We treat both contributions as normally distributed random variables. This way, the fitness of a genotype *g* is given by

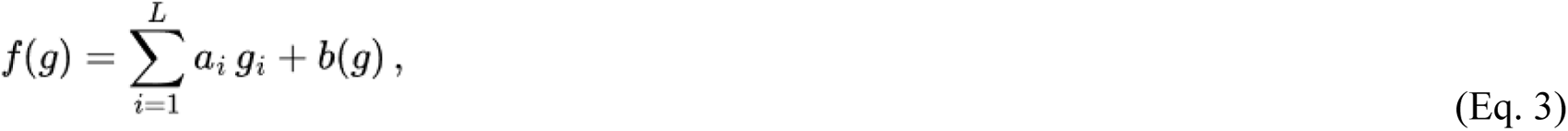

where 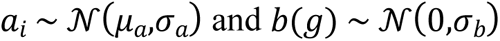; i.e., the additive contributions *a*_*i*_ of each locus *i* follow a normal distribution with mean μ_*a*_ and standard deviation σ_*a*_ and the epistatic components *b*(*g*) follow a normal distribution with zero mean and standard deviation σ_*b*_. Each allele *g*_*i*_ takes the values one or zero, indicating the state of locus *i* in genotype *g*, with a value of one denoting a resistance mutation present in the genotype and the value of zero marking the absence of a resistance mutation at that locus.

This model produces a fitness effect *s*_*j*_(*i*) of a mutation *i* in a background *j* given by

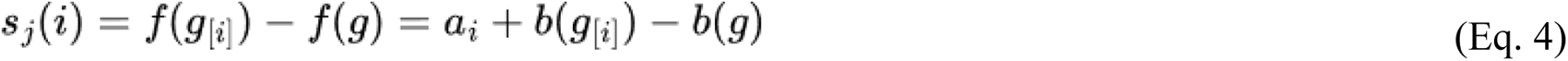

where *g* is the genotype of the background *j* and *g*_[*i*]_ the genotype of the background with the mutation in locus *i*.

To compare the model’s predictions to the data, we constructed a fine grid of pairs of σ_*a*_ and σ_*b*_, for σ_*a*/*b*_ ∈ [0,0.24] with intervals of 0.002. We assumed the mean fitness effect in the model, represented by the parameter μ_*a*_, to be equal to the mean of the experimentally measured fitness effects. For each of these pairs, we generated 10^6^ instances of the model and calculated three summary statistics of the landscape: the mean of fitness effects, the variance of fitness effects, and the gamma epistasis parameters. We used this generated data to obtain the distribution of each summary statistic.

With these distributions, we estimate the likelihood of obtaining the data statistics given the set of model parameters for each summary statistic. In the plots, the likelihood values are represented relative to the maximum likelihood, so all log-likelihoods shown have a maximum of 0.

To compute the experimental data statistics, we used a parametric bootstrap assuming that the fitness effects’ replicate values are normally distributed, taking the mean and standard variation of the replicates as the normal distribution parameters. We also used an alternative bootstrap strategy that produced no qualitative changes in the results, as a test to the assumption’s robustness.

### Analytical estimates of the parameters

A simpler approach is matching the statistics’ mean values directly to the model’s parameters. This method is possible to calculate analytically but less statistically robust since it does not consider the variation among replicates. From Eq. 4, we can see that the mean γ of the fitness distribution and the corresponding variance 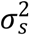 are given respectively by

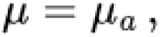

and

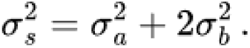

The mean γ parameter is approximated for RMF model as [51]

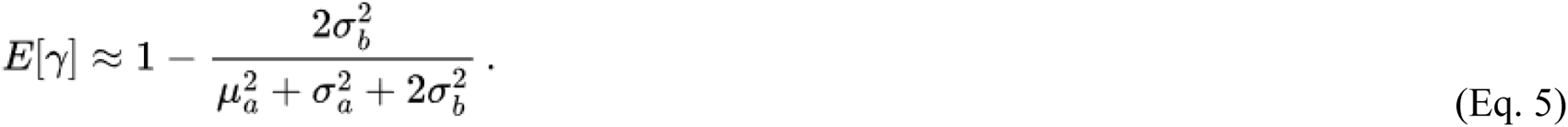

Equating these values to their experimental means 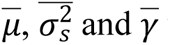 we can solve the system and find

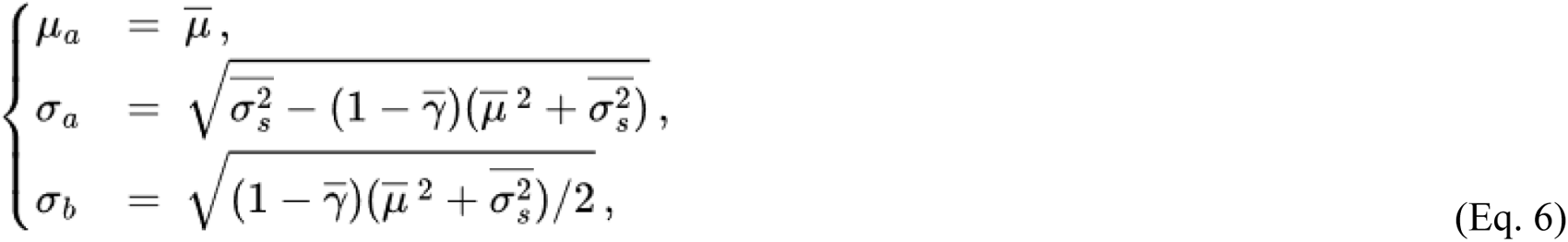

which corresponds to an estimate of the best model parameters based only on the experimental mean values of the statistics. These provide good estimates for the best parameters for the case of single environment landscapes, but fail to describe multiple environment landscapes because the distribution of the statistics is not well approximated by their mean value.

## Acknowledgements

This study was funded by a JPI-AMR grant to AW, RK, and CB (Canadian Institutes of Health Research grant no. 150766, FCT grant JPIAMR/0001/2016). AA was supported by ERC Starting Grant 804569 (FIT2GO) to CB. CB acknowledges support by SNF project grant 315230_204838/1 (MiCo4Sys).

